# Characterization of a fiber-coupled EvenField illumination system for fluorescence microscopy

**DOI:** 10.1101/2021.04.20.440683

**Authors:** Kyla Berry, Mike Taormina, Zoe Maltzer, Kristen Turner, Melissa Gorham, Thuc Nguyen, Robert Serafin, Philip R Nicovich

**Affiliations:** Allen Institute, 615 Westlake Ave N, Seattle, WA 98109, USA; Cajal Neuroscience, 1616 Eastlake Ave E, Ste 300, Seattle, WA 98102, USA

## Abstract

Fluorescence microscopy benefits from spatially and temporally homogeneous illumination with illumination area matched to the shape and size of the camera sensor. Fiber-coupled illumination schemes have the added benefit of straightforward and robust alignment and ease of installation compared to free-space coupled illumination. Commercial and open-source fiber-coupled, homogenized illumination schemes have recently become available to the public; however, there have been no published comparisons of speckle reduction schemes to date. We characterize three different multimode fibers in combination with two laser speckle reduction devices and compare spatial and temporal profiles to a commercial unit. This work yields a new design, the EvenField Illuminator, which is freely available along for researchers to integrate into their own imaging systems.

## 1. Introduction

The ideal characteristics of an illumination system for camera-based fluorescence microscopy are a high degree of spatial and temporal homogeneity, closely matched to the field of view of the sensor, and capable of delivering the required power across the necessary range of wavelengths. Even if corrected computationally, uneven illumination rarely provides satisfactory performance over tiled fields of view or for quantitative analysis. Fluorophore photophysical dynamics in the center versus edges of the field will differ, especially confounding single molecule localization microscopy (SMLM) acquisitions [1–5]

Diode laser sources have become a preferred choice in many fluorescence microscopy systems as their high spatial intensity and monochromaticity provide increased fluorescence emission and multiplexing capabilities. Multiple laser sources can be spatially overlapped into a single multicolor beam can then be easily coupled into a microscope[4,6]. Single-mode diode lasers, either free-space coupled or combined into a single-mode optical fiber, typically output a radially symmetric Gaussian TEM00 mode. This Gaussian illumination profile results in uneven illumination (Fig 1A and B) and must be corrected to provide the desired spatially homogeneous profile (Fig 1C-E). Such a profile can result from passing a Gaussian beam through engineered diffusers, diffractive elements, or refractive beam shapers [1–3,7,8].

**Fig. 1.**
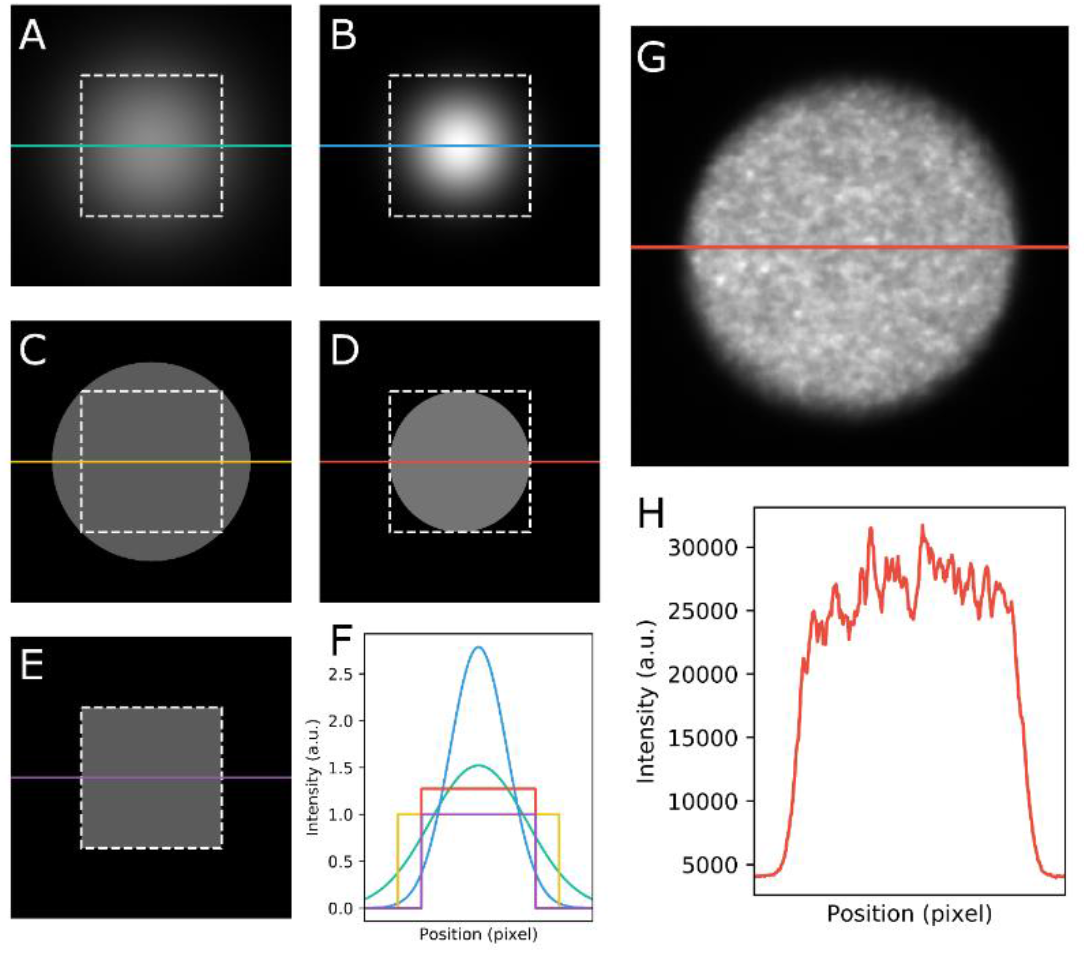
Simulated Gaussian illumination profiles with a standard camera sensor (white dashed box) surrounding FWHM (A) and 1/e^2^ bounding boxes (B). Simulated circular illumination field top hat profiles with camera sensor circumscribed (C) and inscribed (D). Simulated illumination profile of ideal, sensor-matched, square, top hat profile (E). Line profiles of simulated illumination fields with intensities scaled to 1 a.u./pixel for the area within the white box (F). Illumination profile of imaged circular, multi-mode fiber with apparent laser speckle noise (G) and plotted line profile (H).

An alternative approach is to couple the spatially overlapped beams into a multimode optical fiber and project the front face of the fiber onto image plane. Propagating the beam through the multimode fiber results in the original Gaussian shape of the beam to be evenly distributed throughout the fiber core. This results in a more even output profile compared to the Gaussian shape produced from single-mode fibers. Coupling into multimode fibers can be efficient and with high-power multimode laser diodes the output of the illumination system can be sufficient for demanding SMLM applications [3]. Fiber-coupled schemes benefit from straightforward and robust alignment with flexible installation and a small footprint compared to free space schemes.

As multimodal laser light propagates through the fiber, interference effects give rise to the laser speckle phenomenon (Fig 1G, H)[9]. Solutions to mitigate laser speckle often involve “scrambling” the beam to average out the speckle distribution over the time exposure of the camera. Approaches include vibrating the fiber [4], incorporating modulated electro-optical polymers [3,4] or spinning diffusers [1,2,10] in the beam path prior to coupling into a multimode fiber. Notably, these dynamic elements remove the temporal coherence of the illumination light and thus must operate on a timescale much shorter than the desired acquisition. The acquisition rate of a modern sCMOS camera can exceed 300 frames per second over a 2048 × 2048 pixel area, meaning an effective despeckle mechanism should support exposure times of a few milliseconds or shorter.

A circular illumination profile such as from a typical circular-core fiber is difficult to match to a traditional square camera sensor (Fig 1C and D). Either the illumination excites a larger field of view than is captured by the camera resulting in additional, unneeded photobleaching of these areas (Fig 1C), or the field of view includes unilluminated areas (Fig 1D). Both cases make tiling difficult and acquisitions inefficient.

Recently, square-core multimode fibers have become commercially available. The combination of a square-core multimode fiber with a speckle reducer provides a laser-based illumination source with even illumination profile and shape matched to the field of view of a camera sensor. Commercial products employing this approach are available (Andor Borealis, Errol Albedo) as well as recently reported open-source implementations [4]; however, a published comparison of these products and design improvements is not available.

An acceptable illumination system for single molecule fluorescence imaging should provide a homogeneous intensity profile that efficiently fills the shape of a camera sensor to provide flat field images with minimal extraneous photobleaching (Fig 1E). Beam conditioning mechanisms should support fast (< 5 ms) exposure times and high (> kW/cm2) intensities over the field of view [5,11]. Ideally this system would be low cost and straightforward to integrate into a microscope.

Here, we test the illumination profile of three multimode fibers and two speckle reduction schemes against a commercial product, the Andor Borealis. The aim is to identify a combination that is cost-efficient, straightforward to implement, and achieves a spatially and temporally homogenous illumination profile that meets or exceeds the commercial standard for widefield fluorescence microscopy. The final implemented design, coined the EvenField Illuminator, meets these criteria at greatly reduced cost compared to commercial implementations. The EvenField Illuminator, in conjunction with high power multimode laser diodes, has the power throughput necessary for SMLM across an entire 220 μm × 220 μm field of view. Design files, analysis code, and an optical alignment guide for the EvenField Illuminator are freely available at https://github.com/AllenInstitute/EvenField for other researchers to replicate the design and use within their own laboratory.

## 2. Results

Characterization of the spatial and temporal homogeneity of the considered illumination systems was accomplished on an imaging assembly consisting of a diode laser source, a speckle reduction device, a multimode optical fiber, coupling optics to project the fiber exit face onto a slide-mounted concentrated dye solution and the subsequent fluorescence emission image to a CMOS camera (Figure 2A). The collimated beam from the laser source focused onto an Optotune laser speckle reducer (LSR), was recollimated, and coupled into one of several optical fibers. The following three multimode (MM) fibers were considered: a 104 μm diameter circular core MM fiber (Circle), a 150 μm × 150 μm square core MM fiber (Small Square), and a 600 μm × 600 μm square core MM fiber (Large Square). The output from these combinations was characterized and compared to the output of the Andor Borealis system as the speckle reduction, fiber coupling optics, and optical fiber portions of the system.

**Figure 2.**
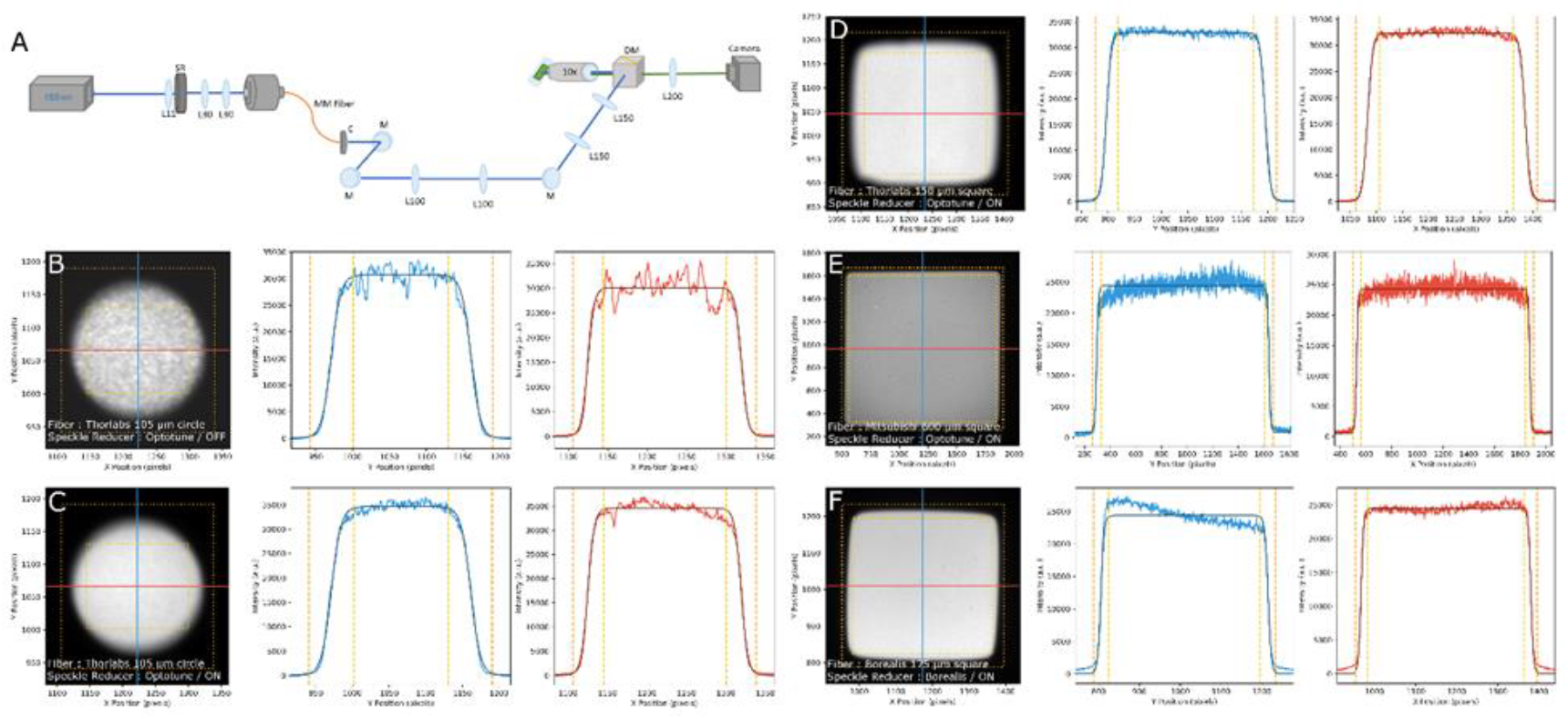
Intensity profiles of imaged multimode fibers (A) Optical diagram used to image fiber faces of scrambled and unscrambled beams. A 488 nm beam is focused onto a laser speckle reducer (SR) and coupled into a multimode fiber (MM). The output beam is then collimated by a 20mm focal length apochromat lens (C) and the fiber face image is focused onto the sample (concentrated fluorescein dye sandwich) using a 10x Nikon objective (NA=0.3). Borealis commercial speckle reduction unit bypasses the SR and is collimated at C. Lxx: Achromatic lens of focal length xx, DM: Dichroic mirror M: Mirror (B-F) Homogeneity images from selected from tested fibers with Optotune or Borealis speckle reducer. Left panel in each shows fluorescence image of projected fiber face on test specimen. Blue and red lines indicate traces along X (red) and Y (blue) axes. Intensity profiles in center and right panel are pulled from X and Y traces (light red or blue), are fit by tanh function (dark red or blue), and plotted. Yellow and orange dashed lines are 99% and 1% levels between the fitted baseline and peak level intensities. These define the inner (yellow) and outer (orange) bounding boxes in intensity images. Tested fibers: Thorlabs 105 μm circular core with Optotune SR off (B) and on (C) (Circle), Thorlabs 150 μm × 150 μm square core fiber with SR on (D) (Small Square), Mitsubishi 600 μm × 600 μm square core fiber with SR on (E) (Large Square), and the commercial Borealis unit with Borealis SR on (F)

Collected images for each of the tested fiber options, with and without speckle reduction, appear in Supplemental Figure 1 with selected images in Figure 2. The calculated centroid of each fiber face in the acquired image and extracted horizontal and vertical line profiles from these images passing through the centroid. The left and right portion of the line profiles are then fit by a hyperbolic tangent function of the form 

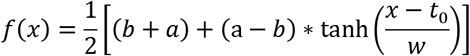

where the baseline background intensity is represented by b, the mean signal intensity of the tophat is represented by a, t0 is the distance from the edge of the image to the break point symmetrically around the origin of spatial coordinate x, and w determines the steepness of the slope. Line profiles of the fiber collected images generally fit well to this function, as shown by the dark fit lines in Figure 2 and Supplemental Figure 1. The values of these fits define bounding boxes at 1% and 99% of peak fit intensity from the collected images, indicated by dashed lines in fiber images of Figures 2 and Supplemental Figure 1.

A qualitative examination of the beam output profiles (Figure 2B-F) show that both the Small Square and Large Square fibers have comparable sensor matching to the Borealis output with the Large Square fiber’s shape most closely matching a square sensor shape. Extracted bounding boxes quantify this fit, with the 10% bounding box representing the extremes of the illumination field and the 90% bounding box representing an optimal area for homogeneous illumination. The normalized difference in area between these two boxes ([1% box area – 99% box area] / [1% box area]) stands as a proxy for the fluorescent emission data that would be lost due to sensor cutoff in an imaging experiment from an imperfect top hat illumination profile, with a small value representing a close match between illumination and sensor.

Consistent with qualitative examination, the Large Square fiber displayed a profile most similar to a top hat with a difference in area of 17.4% compared to 27.6%, 45.1% and 65.5% for the Borealis, Small Square, and Circle fibers, respectively. Image intensity within the 90% bounding box represents 88.0% of the total detected intensity in the Large Square fiber case and speckle reduction active, and this value being 83.7%, 73.3%, 74.2% for the Borealis, Small Square, and Circle fibers. Together these data indicate that the Large Square fiber is the best match to a square sensor form factor.

The homogeneity of illumination was calculated by finding the coefficient of variation (standard deviation divided by mean) of the horizontal, vertical, and 2D intensity profiles found within the 90% bounding box region. The coefficient of variation measured in images at 16 ms exposure time is decreased 1.29− to 2.8-fold (Circle and Small Square, respectively) when the Optotune or Borealis speckle reducer is active. Tested fibers show a coefficient of variation over the 99% bounding box area of 0.0230 (Small Square) to 0.0756 (Circle), with the value for the Borealis (0.059) a small improvement over the Large Square fiber (0.068). Measurements on the XY traces rather than the 2D image show a similar trend, but with the Circle fiber showing the second-best result. This is a consequence of the 2D bounding box being a poor fit for the Circle fiber’s illuminated area. Interestingly, the XY trace of the Borealis data shows an anomalously high value along one axis; this is indicative of the angle-polished fiber face mounted in a component designed for a flat-polished face. Along the other axis the variation of the Borealis illumination is close to that of the Small Square fiber (0.022 for Borealis, 0.020 for Small Square). The values for the tested fibers match favorably with the commercial Borealis unit, with some advantage in homogeneity to the Small Square fiber.

In many experiments, temporal homogeneity of the illumination is as or more important than the spatial homogeneity. The laser speckle reducers are dynamic components so if the timescale of those dynamics is slower than that of the acquisition, the illumination stability could be adversely affected. To test this, we collected a stack of 1000 images for each test fiber with the LSR on and off at various exposure times ranging from 1 ms to 1000 ms. In each frame an average intensity is measured in a 64 × 64 pixel region in the center of the illuminated area, with results appearing in Figure 3A and Supplemental Figure 2. An autocorrelation of these time traces shows the magnitude and timescale of fluctuations in the illumination intensity at different exposure times.

**Figure 3.**
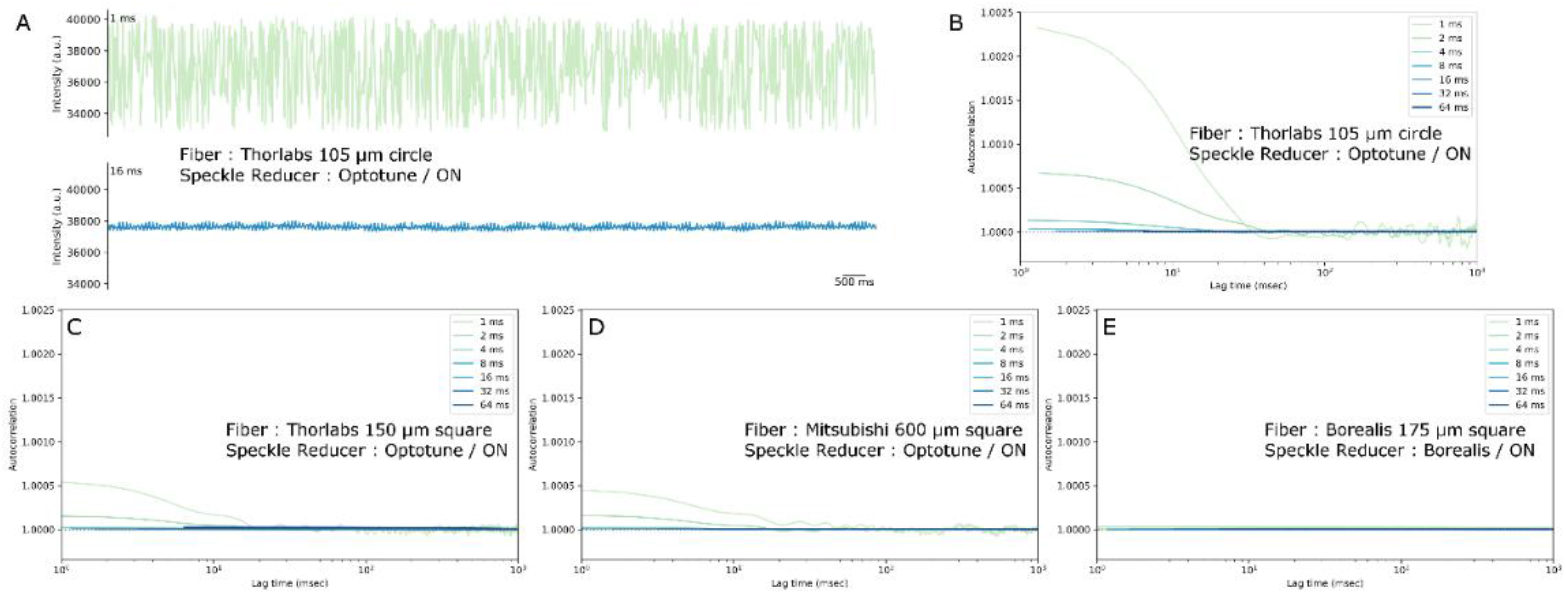
Time domain analysis of speckle reduction in selected tested multimode fibers (A) Example time traces of average 2d intensity profiles at indicated exposure time with Optotune speckle reducer active. Intensity axes are identical for both exposure times. (B-C) Time autocorrelation of average 2d signal intensity profiles taken from indicated fiber at 1, 2, 4, 8, 16, 32, and 64 ms exposure times. All autocorrelation plots are scaled to same x and y axis range for comparison. See Supplemental Figure 2 for plots on independent y axis scales for each panel.

Autocorrelation magnitude analysis (Figure 3B-E) shows large fluctuations apparent at exposure times shorter than 4 ms when using the Optotune LSR. This corresponds well with the nominal 300 Hz oscillation frequency of that device. In contrast, this time dependent speckle reduction was not seen in the Borealis system at the exposure times analyzed suggesting that the Borealis system utilizes a speckle reduction element that operates at a frequency greater than 1000 Hz. Although the Borealis system outperforms the Optotune laser speckle reducer in combination with the tested multimode fibers, the latter speckle reduction scheme is comparable at exposure times > 4 ms with time series coefficient of variations (equivalent to 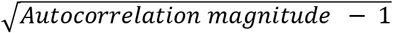) under 0.005 for all tested fibers.

These results confirm that the Optotune LSR with a multimode fiber is a reasonable route to spatially homogeneous illumination. Temporal homogeneity compares well with the commercial standard for exposure times longer than 4 ms. However, the damage tolerance rating for this component is moderate (300 W/cm2, corresponding to μW of power in a μm-scale spot). This is insufficient for adequate excitation over a field of view matching a typical camera sensor. In addition, the inability to perform acquisitions with exposure times < 4 ms is a significant constraint.

To this aim, we created a new, easily deployable, speckle reduction scheme, coined the Evenfield Illuminator. Here, the speckle reduction is accomplished with a rotating 0.5° holographic diffuser placed in the collimated beam path before the fiber input. The diverging beam is collected and focused onto the multimode fiber with an 11 mm focal length aspheric lens (Figure 4A). The fiber position was adjusted using an XYZ translating mount to maximize coupling efficiency into a multimode fiber. With this setup, we achieved coupling efficiencies of 33%-70% across 5 laser lines (405 nm – 750 nm) into the Large Square fiber (Table 1).

**Table 1.** Laser coupling and intensity measurements for the EvenField system using the Large Square fiber and coupled into an Olympus IX83 microscope (Supplemental Figure 5). Maximum output of the two Oxxius lasers exceeds the power capacity of the power meter used, so both the maximum and a reduced power measurement are included.

| Laser Model | Wavelength | Nominal beam quality ($M^2$) | Power at laser exit | Power at fiber output | Power at objective pupil | Coupling efficiency, fiber exit | Coupling efficiency, objective pupil | Intensity at specimen, 220 $\mu\text{m}$ x 220 $\mu\text{m}$ field |
| --- | --- | --- | --- | --- | --- | --- | --- | --- |
| **Vortran Stradus 405-100** | 405 nm | < 1.25 | 99.7 mW | 33.2 mW | 14.02 mW | 33.3% | 14.1% | 0.029 kW/cm<sup>2</sup> |
| **Vortran Stradus 488-150** | 488 nm | < 1.25 | 149.6 mW | 101.3 mW | 72.0 mW | 67.7% | 48.1% | 0.148 kW/cm<sup>2</sup> |
| **Vortran Stradus 561-50** | 561 nm | < 1.1 | 53.0 mW | 37.1 mW | 26.9 mW | 70.0% | 50.7% | 0.056 kW/cm<sup>2</sup> |
| **Oxxius LBX-638 HPE** | 638 nm | 22.4 x 2.2 | 190 mW | 98.0 mW | 86.4 mW | 51.6% | 45.5% | 0.179 kW/cm<sup>2</sup> |
|  |  |  | - | 780 mW | 575 mW | - | - | 1.19 kW/cm<sup>2</sup> |
| **Oxxius LBX-750 HPE** | 750 nm | 59.7 x 4.78 | 307 mW | 134.0 mW | 96.5 mW | 43.6% | 31.4% | 0.020 kW/cm<sup>2</sup> |
|  |  |  | - | 639.8 mW | 492 mW | - | - | 1.02 kW/cm<sup>2</sup> |

**Figure 4.**
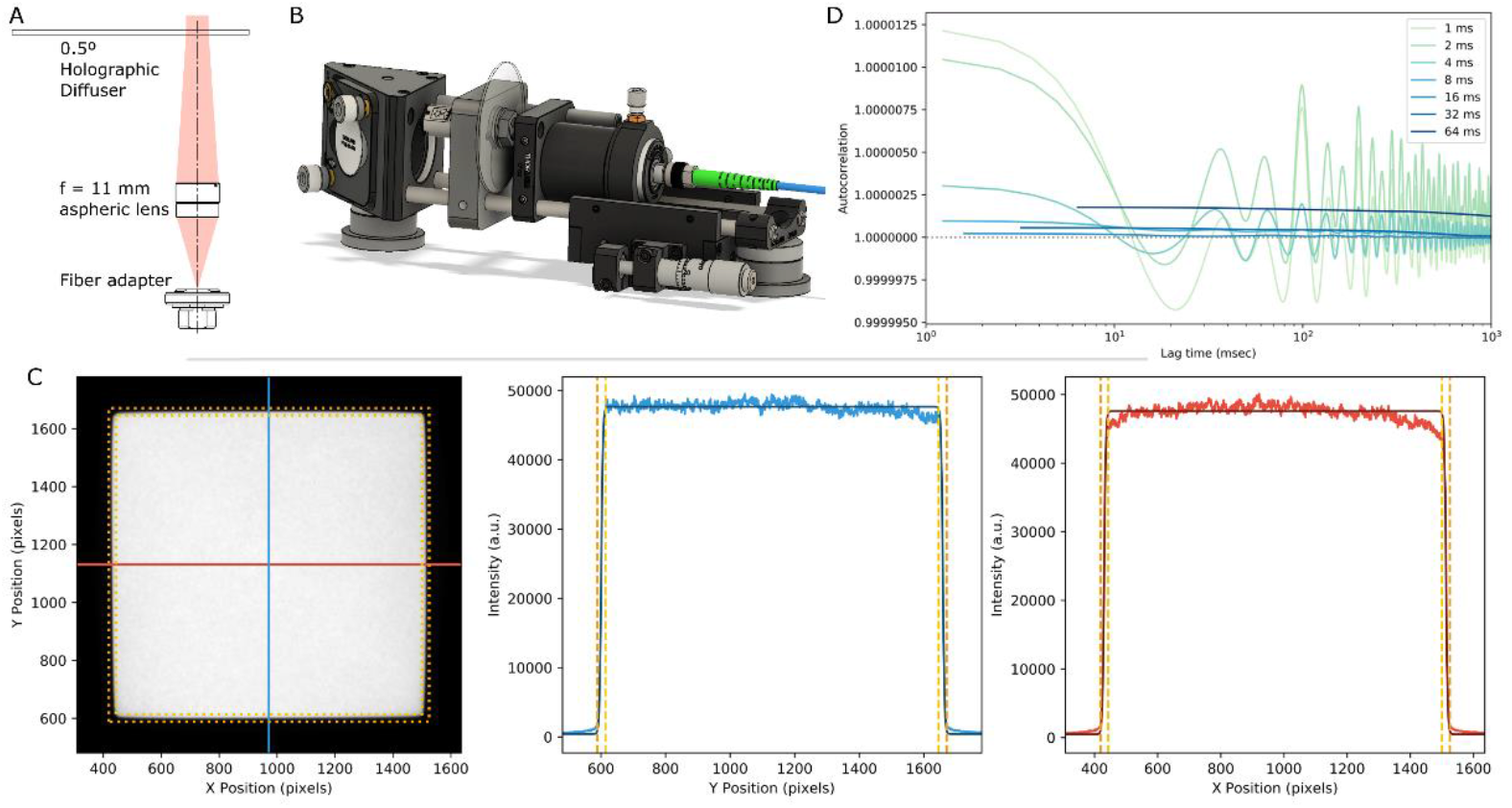
Evenfield illumination scheme for improved temporal and spatial homogeneity and power throughput (A) Optical diagram of new speckle reduction scheme. The collimated laser beam is sent through a 0.5° holographic diffuser and the scrambled diverging beam is focused onto the fiber input. (B) CAD rendering of assembled beam scrambler and fiber coupler (C) Imaged Large Square fiber face (left) and intensity profile traces (middle and right, at indicated lines in left) with rotating diffuser turned on. Dashed lines indicate 10% and 90% bounding regions. (D) Time autocorrelation plots of average 2d intensity profiles from Large Square fiber with rotating diffuser on.

The Large Square fiber demonstrated the best performance with respect to illumination area sensor matching and comparable performance to the other fiber options for spatial and temporal homogeneity measurements. As such, we chose to continue development with the Large Square as the primary option. The Large Square fiber output profile with EvenField Illuminator speckle reduction was then analyzed as before, yielding an excellent top hat profile with a coefficient of variation of 0.0289 for the fiber face image within the 90% bounding region (Figure 4C). The illuminated area shows a similar match to the ideal square form factor as before, with (1% box area – 99% box area)/(1% box area) value of 14.5% and 89.9% of the illumination intensity falling within the 99% bounding box area. These values can be slightly improved by using a 10x/0.3 NA objective to image the exit face of the fiber rather than a FiberPort unit. With this change, the intensity variation falls to 0.0265, box area difference to 8.8%, and illumination within the 99% bounding box rising to 93.8%. While the FiberPort is more convenient to install, the objective shows better performance, including improvements in vignetting and pincushion distortion; the application will dictate which of these solutions is preferred.

Parameters defined by the ISO 13694:2000 [12] and prior work [2] are useful to place the performance of this unit in context. For the Large Square fiber configuration with objective collimator, edge steepness is 0.041, plateau uniformity 0.038, and flatness factor 0.933; with the FiberPort collimator edge steepness is 0.044, plateau uniformity 0.037, and flatness factor 0.845. Both implementations compare favorably to demonstrated schemes for free-space-coupled flat field devices [2,4]. Speckle contrast with the Large Square fiber is 0.018 and 0.019 for the objective and FiberPort collimation configurations, respectively.

Temporal stability is improved with the EvenField illuminator, with the coefficient of variation remaining below 0.36% for all tested exposure times from 1 ms to 1000 ms. Varying the voltage to the EvenField motor confirms that the speckle reduction is driven by this motor rotation and a reduction in voltage (and consequently motor speed) still produces performance matching the commercial standard. Such considerations may be valuable for component stability or alternatively, increasing motor speed being a straightforward route to even further improving stability at short exposure times. Interestingly oscillations in the autocorrelation signal in these measurements show the periodic nature of the speckle reduction method. This contrasts to the Optotune LSR, which is reportedly driven by a random process. The magnitude of the oscillations seen in the EvenField case are small but are worth noting if these periodic fluctuations could be inadvertently enhanced by downstream analysis.

As an additional adaption, we performed the same measurements with the EvenField speckle reducer and the Small Square fiber (Supplemental Figure 6). Spatial and temporal homogeneity are similar to the Large Square case (0.0347 coefficient of variation of intensity over the 99% bounding box, with temporal coefficients of variations below 0.0086 for all exposure times tested). Sensor matching performance is lower relative to the Large Square case, reflecting what is seen with tests using the Optotune LSR (bounding box difference of 38.8% with 77.6% of illumination within the 99% bounding box). Values for edge steepness (0.073), plateau uniformity (0.127), flatness factor (0.909), and speckle contrast (0.022) are similarly relaxed relative to the Large Square implementation, while remaining similar or exceeding other demonstrated flat fielding implementations. The Small Square fiber is an off-the-shelf, low-cost part compared to the Large Square fiber, which is custom-made. When there is room for some compromise in performance matching the area of the camera sensor, this configuration could be advantageous.

As a demonstration of the EvenField system with biological specimen, we performed a tiled image acquisition of a multicolor fluorescently labeled mouse brain section using a commercial microscope platform. The EvenField configuration with the Large Square fiber was coupled into an Olympus IX83 microscope (Supplemental Figure 5). Images of a slide-mounted 25 μm thick mouse brain section labeled with DAPI, AlexaFluor 555, and AlexaFluor 488 were acquired as a 6×12 tile region. Some residual inhomogeneity apparent in the image tiles (Figure 5A) is straightforward to correct with an internal flat-field averaging procedure. The corrected and stitched tiles result in an image with minimal appearance of tiling borders (Figure 5B).

**Figure 5.**
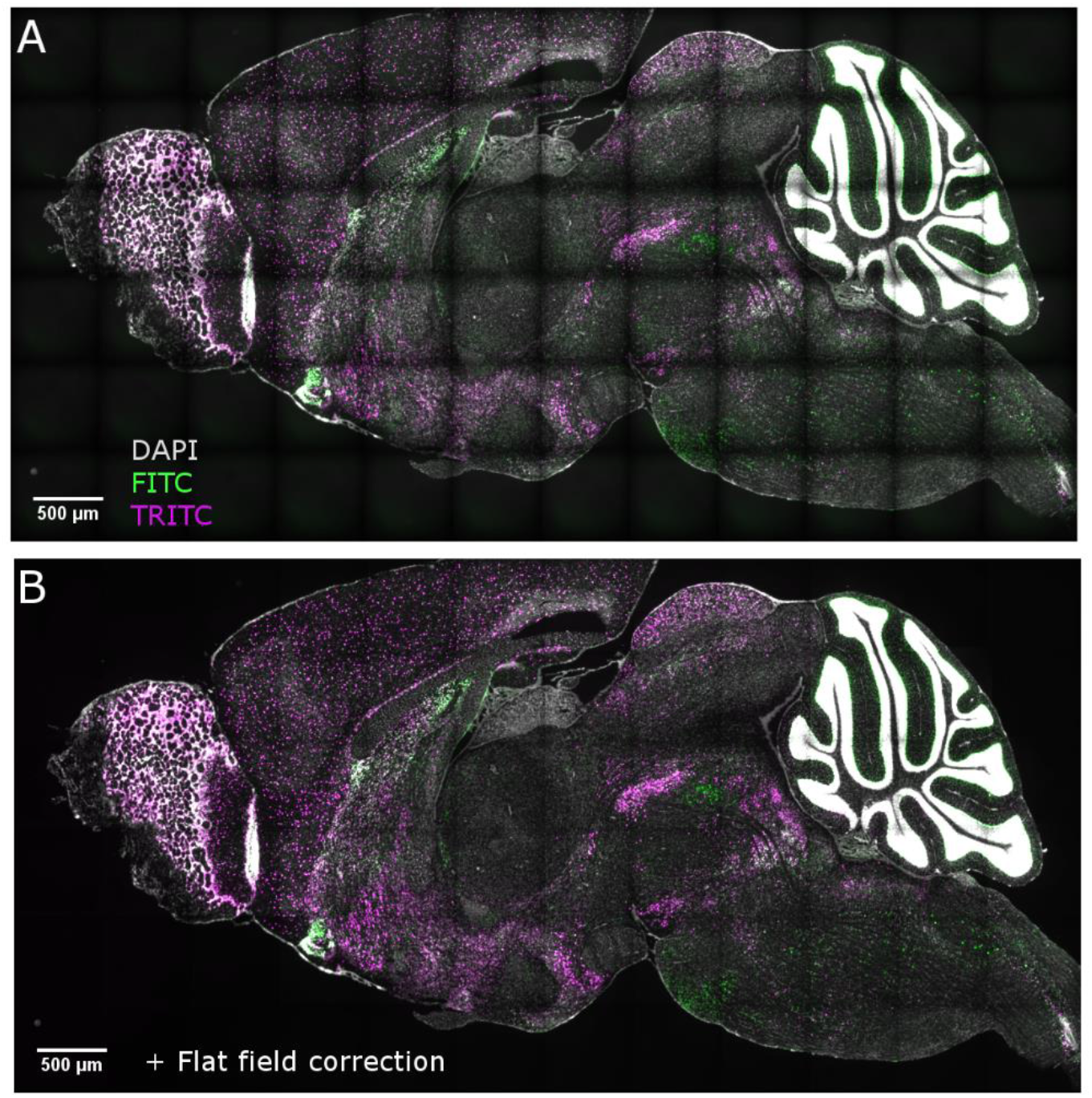
Multi-tile image acquisition with described Evenfield Illumination System A 6×12 tiled 3 channel image of mouse sagittal section was acquired with (B) and without (A) flat-field correction based on background of empty tiles in each channel’s acquisition. Images were acquired on an inverted Olympus microscope with Evenfield illumination coupling is as shown in Supp Fig 5.

This implementation includes two multimode laser diodes with high output powers (Oxxius LBX-638HPE and LBS-750HPE; 1.1 W and 1.2 W nominal output) and a beam quality heavily diverging from a TEM00 mode. With nominal M2 values of 22.2 and 59.7 for the 638 nm and 750 nm unit, respectively, a high coupling efficiency is maintained (51.6% and 43.6%, respectively). Maximum output power of these units delivers 780 mW or 640 mW to the fiber exit and 575 mW or 492 mW to the specimen, for the 638 nm or 750 nm units, respectively. This corresponds to 1.19 kW/cm2 or 1.02 kW/cm2 over a 220 μm × 220 μm area (i.e. a typical FOV for a 4.2 MP sCMOS camera with 60x magnification). This value meets the nominal requirements to drive fluorophore photophysics in SMLM.

## 3. Discussion and Conclusion

We characterize the performance of multimode fibers with laser speckle reducers for the application of fluorescence microscopy. Both spatial and temporal homogeneity is considered, as well as the sensor-matching performance of the illumination. A commercial unit acts as the standard to benchmark tested systems. Results show that the best sensor-matching performance is achieved with a 600 μm × 600 μm square-core custom made for this application. With the commercially available Optotune LSR, spatial homogeneity is sufficient but temporal homogeneity is lacking for exposure times < 4 ms and rated laser powers are insufficient. We designed and implemented the EvenField laser speckle reducer to mitigate these shortcomings.

The EvenField Illuminator demonstrates to produce a homogenous, square, speckle free, illumination profile meeting or exceeding the quality of standard commercial systems or other demonstrated approaches for multi-color, multi-tile, widefield fluorescence imaging. With a large core multimode fiber included in the design, high coupling efficiencies can be achieved. In addition, the design allows for the use of high power multimode laser diodes, providing an illumination system for fiber-coupled, sensor-matched illumination over the entirety of a 4.2 MP sCMOS camera at 60x magnification.

The design can be easily implemented into commercial and open source laser diode combiners (such as Nicolase 3600) for less than $3500 USD. A full bill of materials, CAD files, and an optical alignment guide are provided through the publicly available EvenField GitHub repository hosted by the Allen Institute (https://github.com/AllenInstitute/EvenField). The repository provides all Python analysis code utilized in this manuscript for researchers to use and evaluate the field flatness of their illumination systems.

Together the EvenField system provides a straightforward, low-cost design for sensor-matched, flat-field illumination for fluorescence microscopy. The power density delivered is sufficient for SMLM across large fields of view while maintaining the flexibility and robustness of a fiber-coupled laser illumination design.

## 4. Methods

### Dye Sample Preparation for Fiber Profile Experiments

A filtered, concentrated fluorescein dye solution was pipetted into a solution chamber created by sandwiching double sided tape between a glass slide and coverslip. The dye chamber was then sealed with nail polish and mounted orthogonally to the illumination objective. Dye solution was concentrated enough to show self-quenching properties of fluorescein to minimize imaging volume.

### Optotune Beam homogenizer

A 488 nm beam was focused onto a laser speckle reducer (Optotune LSR-3005-17S-VIS) by a 11 mm focal length aspheric lens. The scattered light was then collected, recollimated, and focused by a 30-30mm matched achromatic lens pair into one of three multimode fibers mounted onto a xyz translating stage as described in Figure 2A. Fiber position was adjusted until efficient coupling was achieved (>50% power throughput).

### Fiber illumination field profile imaging

Three different fiber illumination profiles were tested and compared to a commercial Borealis unit. A custom, multimode square (MM) core fiber (600 μm × 600 μm core size, 0.15 NA, P/N: KV95P2 ST600SQH) from Mitsubishi Cable Industries, LTD. (MCI) was collimated using an aspheric lens system (Thorlabs PAF2P-18A) or 10x/0.3 NA objective (Nikon N10X-PF). The collimated beam is relayed to the image plane through two pairs of 1:1 relay telescopes, first through a pair of f = 100 mm achromatic doublets, then a pair of f = 150 mm achromatic doublets, then through an excitation filter (Semrock FF01-475/35) and reflected by a dichroic mirror (Thorlabs MD498) to a 10x/0.3 NA objective (Nikon N10X-PF) to the sample.

Emission is collected by the same objective, passed through the dichroic and emission filter (Semrock FF01-530/43), before focusing through a tube lens (f = 200 mm achromatic doublet) to detected with CMOS camera (Chameleon3 CM3-U3-50S5M for Optotune LSR testing and Hamamatsu C11440-22CU for EvenField testing). Data acquisition was performed in Micromanager. Similarly, a 105 μm core diameter MM fiber (Thorlabs M43L05), 150 μm × 150 μm square core MM fiber (Thorlabs M103L05), and the Borealis commercial unit were imaged. All fiber faces were imaged with the speckle reducer on and off at 1, 2, 4, 8, 16, 32, 64, 125, 250, 500, and 1000 ms exposure times. Laser powers were adjusted at each exposure time to ensure no oversaturation of pixels.

### Fiber profile analysis

Fiber profiles were analyzed with custom Python code that first subtracts off a dark field image from the acquired fiber image. After, the fiber illumination profile was identified in the image by Otsu thresholding the fiber image from the background and the image is binarized so that the centroid coordinates of the fiber face can be identified. Using these centroid coordinates, horizontal and vertical line profiles are extracted from the image and their intensities were plotted against the image pixel coordinates.

Line profiles were then fit by a Tanh function using Nelder-Mead optimization. Tanh fits were plotted on top of actual line profiles in a darker shade. Bounding boxes were identified from fits at 1% and 99% intensities and plotted as orange and yellow dashed lines, respectively. Coefficient of variation was calculated as the variation in intensity of pixels along horizontal, vertical, and 2D profiles above within the 99% bounding box.

ISO 13694:2000 [12] parameters were calculated following [2]. Edge regions at 10% and 90% intensity are identified from intensity values from the centroid line traces. Edge steepness is defined as (width of 10% area – width of 90% area) / (width of 10% area). Flatness factor is defined as the standard deviation of the intensity within the 99% bounding box normalized to the mean intensity in this area. Plateau uniformity is defined as the FWHM of the peak intensity, normalized to the maximum intensity, within the 90% bounding box.

Speckle contrast is defined following [4] as the standard deviation of the centroid-intersecting intensity line trace minus the tanh fit line, normalized to the mean of the fit line value. Values here are restricted to those falling within the 99% bounding box. Individual values for the X and Y trace are calculated and the mean taken as the final value.

Analysis results are collected into a JSON file and saved to disk. Analysis code can be found in the EvenField GitHub repo (https://github.com/AllenInstitute/EvenField/tree/master/Analysis/Code).

### Measuring speckle reduction timescale

Speckle timescale was measured by imaging a small region located in the center of the fiber image and acquiring 1000 frames at the maximum interval rate the camera allowed for 1, 2, 4, 8, 16, 32, and 64 ms exposure times. This was performed individually for each fiber analyzed and the commercial Borealis unit. Measurements were then analyzed using a custom Python script that first crops the image to a 64×64 pixel region at the image center. Then, it calculates the average intensity of each frame, uses camera metadata to create timestamps for each measurement, and saves a plot of the average intensity over time at each exposure time for each fiber. With this information, the script then performs a time series autocorrelation of the average intensity over time and plots each data set as its own line on a graph based on exposure time.

Analysis results are saved to a separate JSON file. Analysis code can be found in the EvenField GitHub repo above.

### Evenfield Illuminator Speckle Reduction

A 50mm diameter, 0.5°, holographic diffuser (Edmund Optics #47-989) was removed from its mount and was attached to a miniature gear motor (Pololu #3036) through a custom milled mounting assembly (see associated CAD). A DC power supply was used to power the motor with 12V being supplied apart from minimum required voltage testing experiments (Supplemental Figure 4). A collimated 488 nm laser was then sent through the rotating diffuser and the resulting diverging beam was focused onto one of two square core multimode fibers by a 11 mm focal length aspheric lens (Thorlabs C220TMD-A). The position of the fiber with respect to the aspheric lens was then adjusted using an XYZ mount (Thorlabs CT1 and CT102) until maximal coupling efficiency was achieved. Voltage to the rotating diffuser motor was provided with a variable power supply. For motor speed experiments this voltage supplied to the motor was varied.

A full parts list for the EvenField Illuminator is located in the EvenField GitHub repo (https://github.com/AllenInstitute/EvenField/blob/master/Hardware).

### EvenField Fluorescent Imaging

The Evenfield Illuminator Speckle Reduction system (Figure 4A) was placed in the output beam path of a laser diode combiner[6] (Nicolase 3600) and coupled into the custom, multimode square (MM) core fiber (600 μm × 600 μm core size, 0.15 NA) from Mitsubishi Cable Industries. The output beam from the MM fiber was then collimated using an 18.4 mm focal length aspheric FiberPort collimator (Thorlabs PAF2P-18A) and aligned into an Olympus IX83 microscope (Supplemental Figure 5).

A 25 μm whole mouse sagittal section stained by double fluorescent in-situ hybridization previously prepared by the Allen Institute as described previously[13] was repurposed for imaging. In brief, fresh frozen tissue sections were collected and fixed onto glass slides. Then, Gad1 and tdTomato riboprobes labeled with digoxigenin (DIG) and dinitrophynl-11-UTP (DNP), respectively, were simultaneously hybridized to the sample. An amplified signal for each probe was then sequentially generated with tyramide (anti-DIG-HRP + tyramide-biotin followed by anti-DNP-HRP + tyramide-DNP) and labeled with streptavidin-Alexa-Fluor 488 and anti-DNP-Alexa-Fluor 555, respectively. DAPI staining was subsequently preformed prior to coverslipping.

A 6×12 tile image of the sample was taken in three colors at 10x magnification (Olympus UPLSAPO10X2) with sequential excitation using 405, 488, and 568 nm Vortran Stradus laser diodes with 5% overlap. Camera exposure time was set to 150 ms for all samples. Emitted fluorescence was separated from the excitation lasers using Semrock FF01-452/45, FF01-525/45, and FF01-609/54 emission filters for the DAPI, AlexaFluor 488 and AlexaFluor 555 channels, respectively, and captured with a CMOS camera (Hamamatsu C13440-20CU). A series of images at 6 planes were taken at each tile location with z step size of 1 μm. Maximum intensity projections of the tile stacks were output by the imaging software and saved as TIFF files along with pixel coordinates for individual MIPs.

## Funding

### Acknowledgments

The authors thank Brian Long from the Allen Institute for recommendations with optical alignments for tiled imaging experiments. In addition, the authors would like to dedicate this paper to the vision, encouragement, and long-term support of our founder, Paul G. Allen.

### Disclosures

The authors declare no conflicts of interest.

### Data availability

Data underlying the results presented in this paper are not publicly available at this time but may be obtained from the authors upon reasonable request.

## Notes

### Competing Interest Statement

The authors have declared no competing interest.

## References

1. K. M. Douglass, C. Sieben, A. Archetti, A. Lambert, and S. Manley, “Super-resolution imaging of multiple cells by optimized flat-field epi-illumination,” Nat. Photonics 10(11), 705–708 (2016).

2. S. Manley, K. Ibrahim, and D. Mahecic, “Characterization of flat-fielding systems for quantitative microscopy,” Opt. Express (2020).

3. J. Deschamps, A. Rowald, and J. Ries, “Efficient homogeneous illumination and optical sectioning for quantitative single-molecule localization microscopy,” Opt. Express 24(24), 28080 (2016).

4. D. Schröder, J. Deschamps, A. Dasgupta, U. Matti, and J. Ries, “Cost-efficient open source laser engine for microscopy,” Biomed. Opt. Express 11(2), 609 (2020).

5. K. Kwakwa, A. Savell, T. Davies, I. Munro, S. Parrinello, M. A. Purbhoo, C. Dunsby, M. A. A. Neil, and P. M. W. French, “easySTORM: a robust, lower-cost approach to localisation and TIRF microscopy,” J. Biophotonics 9(9), 948–957 (2016).

6. P. R. Nicovich, J. Walsh, T. Böcking, and K. Gaus, “NicoLase - An open-source diode laser combiner, fiber launch, and sequencing controller for fluorescence microscopy,” PLoS One 12(3), (2017).

7. F. Stehr, J. Stein, F. Schueder, P. Schwille, and R. Jungmann, “Flat-top TIRF illumination boosts DNA-PAINT imaging and quantification,” Nat. Commun. 10(1), (2019).

8. S. Hwang, T. Kim, J. Lee, and T. J. Yu, “Design of square-shaped beam homogenizer for petawatt-class Ti:sapphire amplifier,” Opt. Express 25(9), 9511 (2017).

9. W. Osten, “Dainty, J. C. (ed.), Laser Speckle and Related Phenomena. 2nd enlarged ed. Berlin-Heidelberg-New York-Tokyo, Springer-Verlag 1984. XVII, 342 S., 146 Abb., DM 75,—. US $ 29.40. ISBN 3-540-13169-8 (Topics in Applied Physics 9),” ZAMM - J. Appl. Math. Mech. / Zeitschrift für Angew. Math. und Mech. 65(9), 459–459 (1985).

10. T. Stangner, H. Zhang, T. Dahlberg, K. Wiklund, and M. Andersson, “Step-by-step guide to reduce spatial coherence of laser light using a rotating ground glass diffuser,” Appl. Opt. 56(19), 5427 (2017).

11. S. Van De Linde, R. Kasper, M. Heilemann, and M. Sauer, “Photoswitching microscopy with standard fluorophores,” Appl. Phys. B Lasers Opt. 93(4), 725–731 (2008).

12. ISO, “ISO 13694:2018, Optics and photonics — Lasers and laser-related equipment — Test methods for laser beam power (energy) density distribution,” (2018).

13. C. L. Thompson, S. D. Pathak, A. Jeromin, L. L. Ng, C. R. MacPherson, M. T. Mortrud, A. Cusick, Z. L. Riley, S. M. Sunkin, A. Bernard, R. B. Puchalski, F. H. Gage, A. R. Jones, V. B. Bajic, M. J. Hawrylycz, and E. S. Lein, “Genomic Anatomy of the Hippocampus,” Neuron 60(6), 1010–1021 (2008).

