## Supplemental Figures for "Characterization of a fiber-coupled EvenField illumination system for fluorescence microscopy"

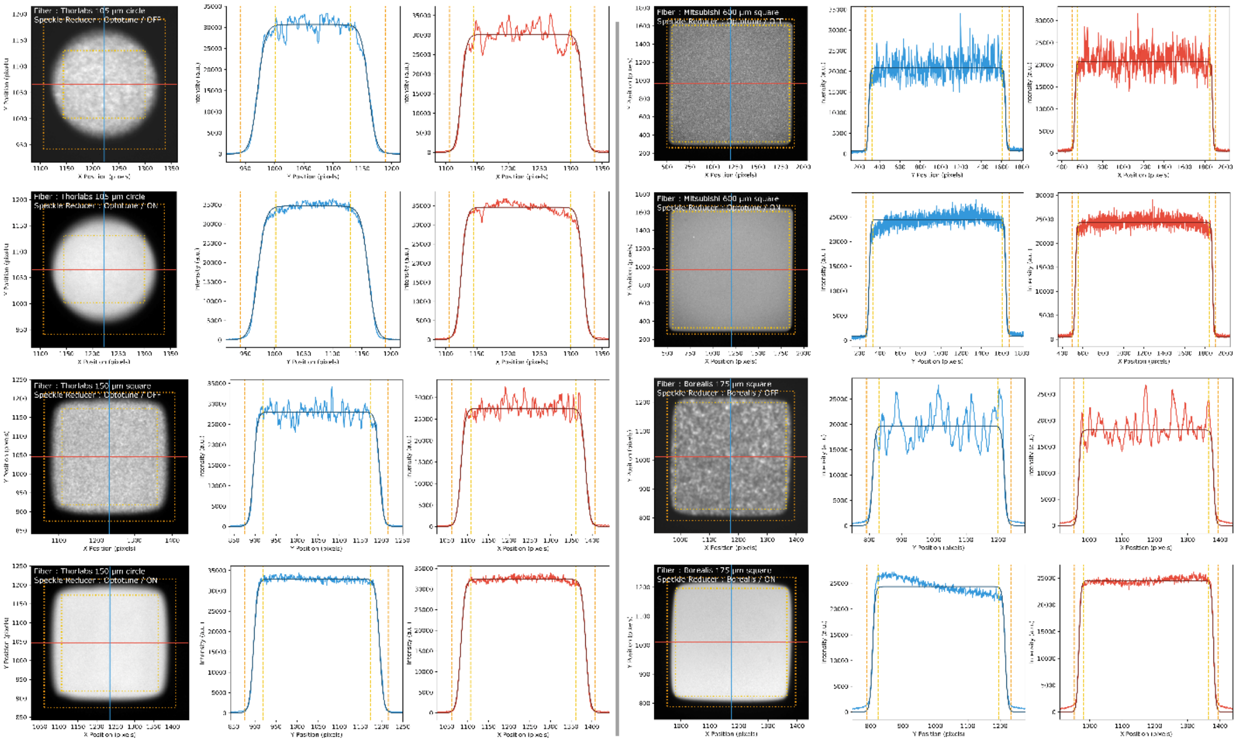


Figure S1 – Homogeneity images from all fibers using Optotune or Borealis speckle reducer. Left panel in each shows fluorescence image of projected fiber face on test specimen. Blue and red lines indicate traces along X (red) and Y (blue) axes. Intensity profiles in center and right panel are pulled from X and Y traces (light red or blue), are fit by tanh function (dark red or blue), and plotted. Yellow and orange dashed lines are 99% and 1% levels between the fitted baseline and peak level intensities. These define the inner (yellow) and outer (orange) bounding boxes in intensity images. Each fiber is shown twice, once with speckle reducer inactive (upper of pair) and with speckle reducer active (lower of pair).

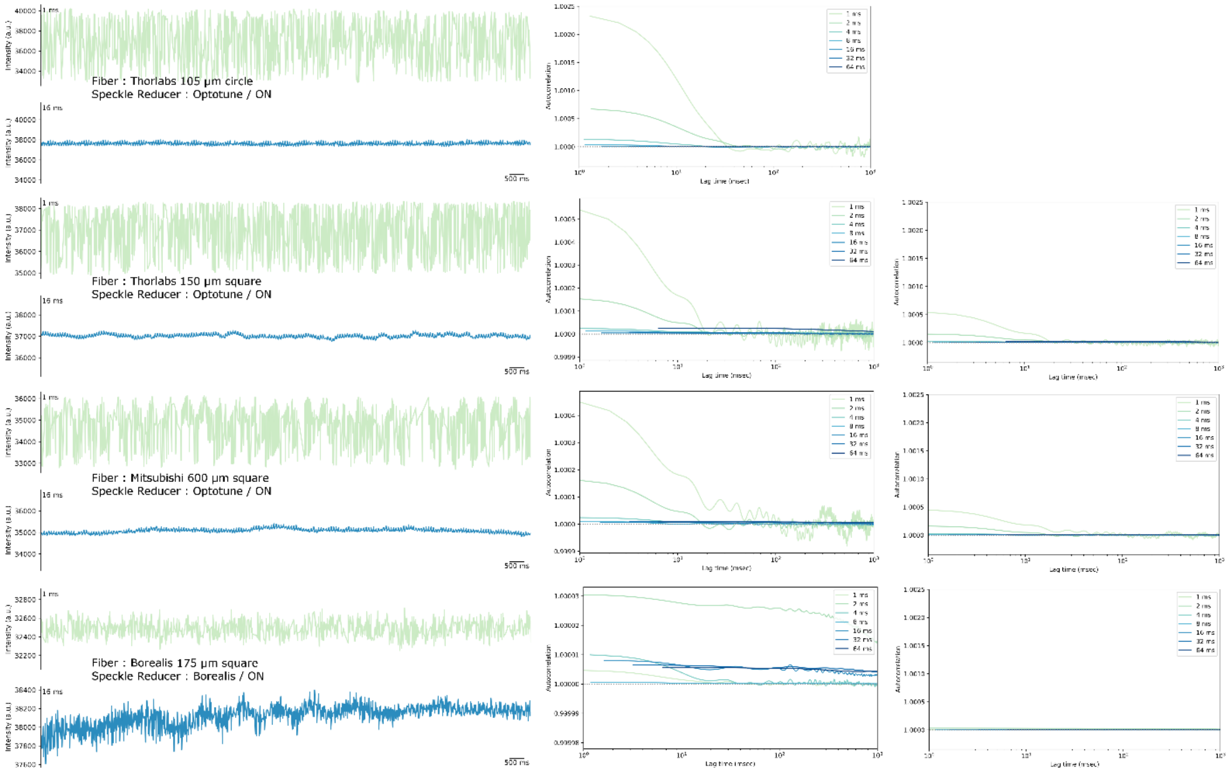


Figure S2 - Time domain analysis of speckle reduction in all tested multimode fibers.

Left column – time traces of average 2d intensity profiles at indicated exposure time with Optotune speckle reducer active. Intensity axes are identical for both exposure times shown.

Center column - Time autocorrelation of average 2d signal intensity profiles taken from indicated fiber at 1, 2, 4, 8, 16, 32, and 64 ms exposure times. Axes scales are independent between rows.

Right column – Data as in center column, but with axes scales matching that of the circular fiber in top row.

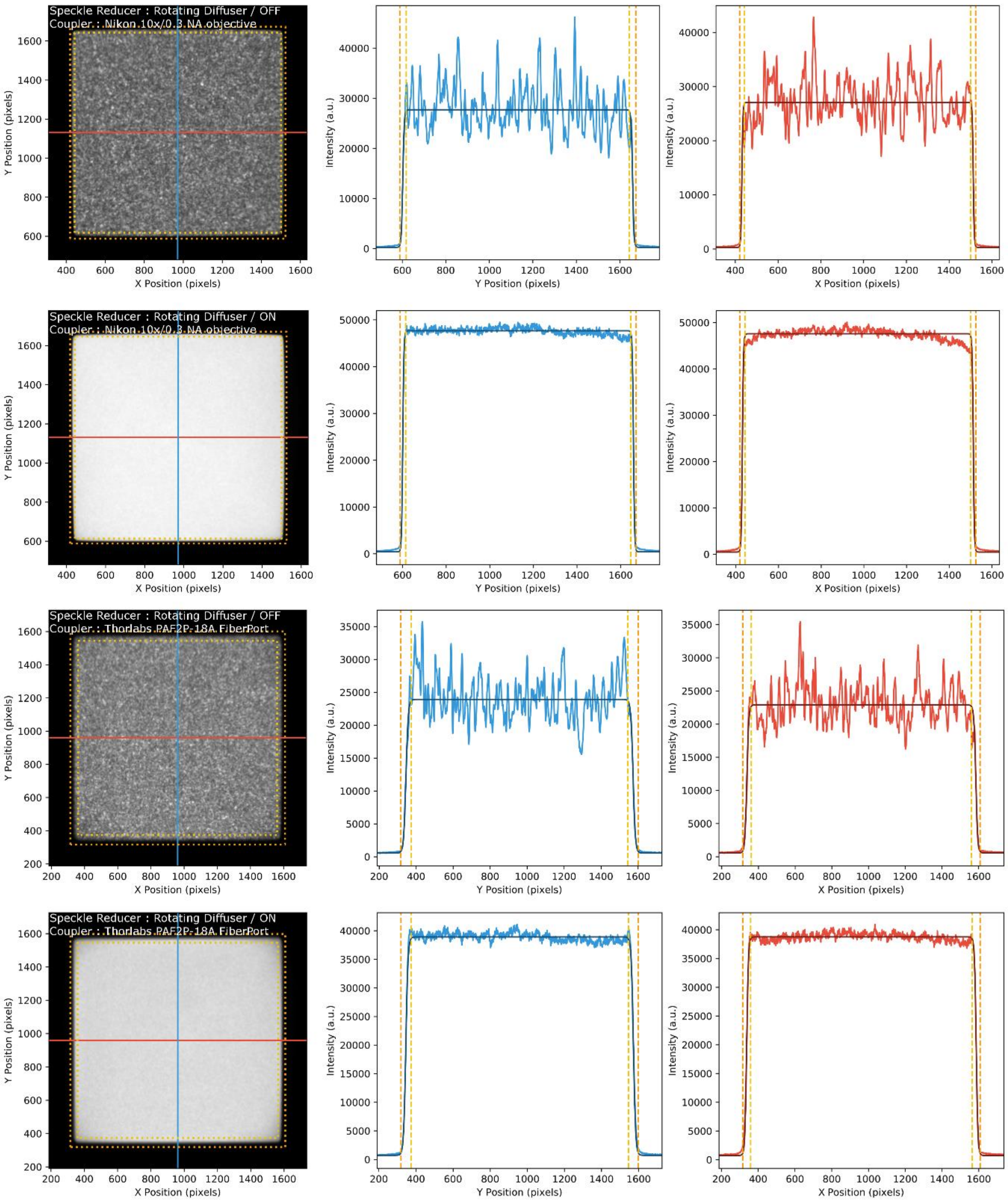


Figure S3 – Homogeneity images from EvenField speckle reducer with Large Square fiber and with objective (upper pair) or FiberPort (lower pair) collimator at fiber exit. Left panel in each row shows fluorescence image of projected fiber face on test specimen. Blue and red lines indicate traces along X (red) and Y (blue) axes, with corresponding intensity profiles and fits in center and right columns. Dashed lines are 99% and 1% levels, defining the inner (yellow) and outer (orange) bounding boxes in intensity images. Each combination is shown twice, once with speckle reducer inactive (upper of pair) and with speckle reducer active (lower of pair).

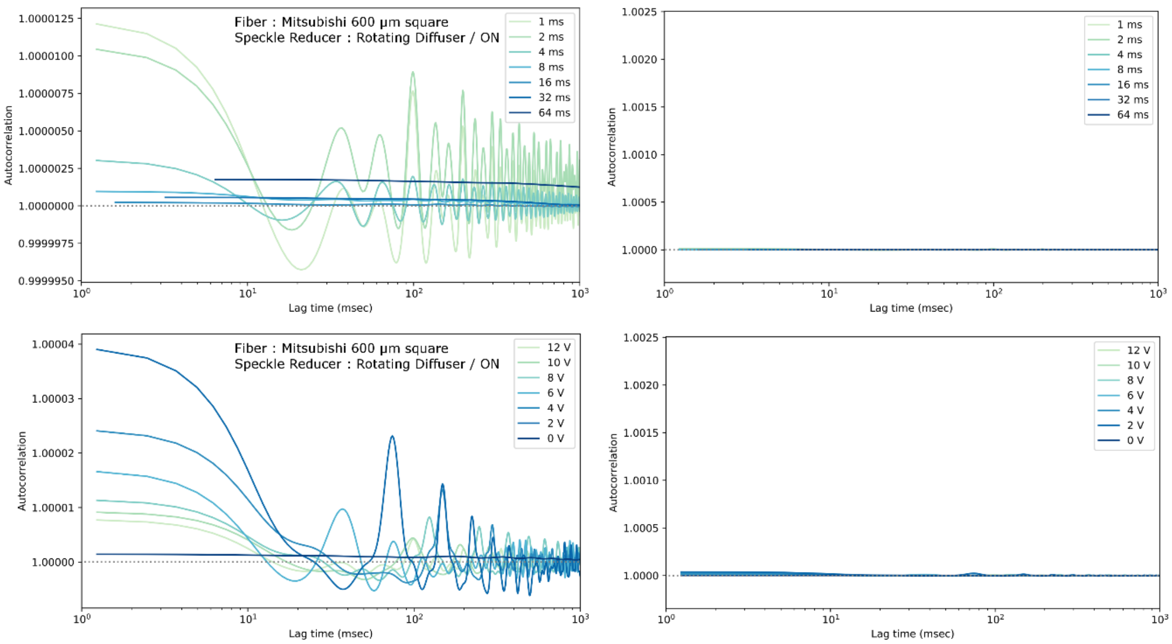


Figure S4 - Time domain analysis of speckle reduction with EvenField speckle reducer.

Top row - Time autocorrelation of average 2d signal intensity profiles taken from Large Square fiber at 1, 2, 4, 8, 16, 32, and 64 ms exposure times. Axes scaled to emphasize these data (left) and scaled to match that of circular fiber with Optotune speckle reducer (Figure 3 and Supplemental Figure 2).

Top row - Time autocorrelation of average 2d signal intensity profiles taken from Large Square fiber at varying voltage to motor and 1 ms exposure time. Axes scaled to emphasize these data (left) and scaled to match that of circular fiber with Optotune speckle reducer (Figure 3 and Supplemental Figure 2).

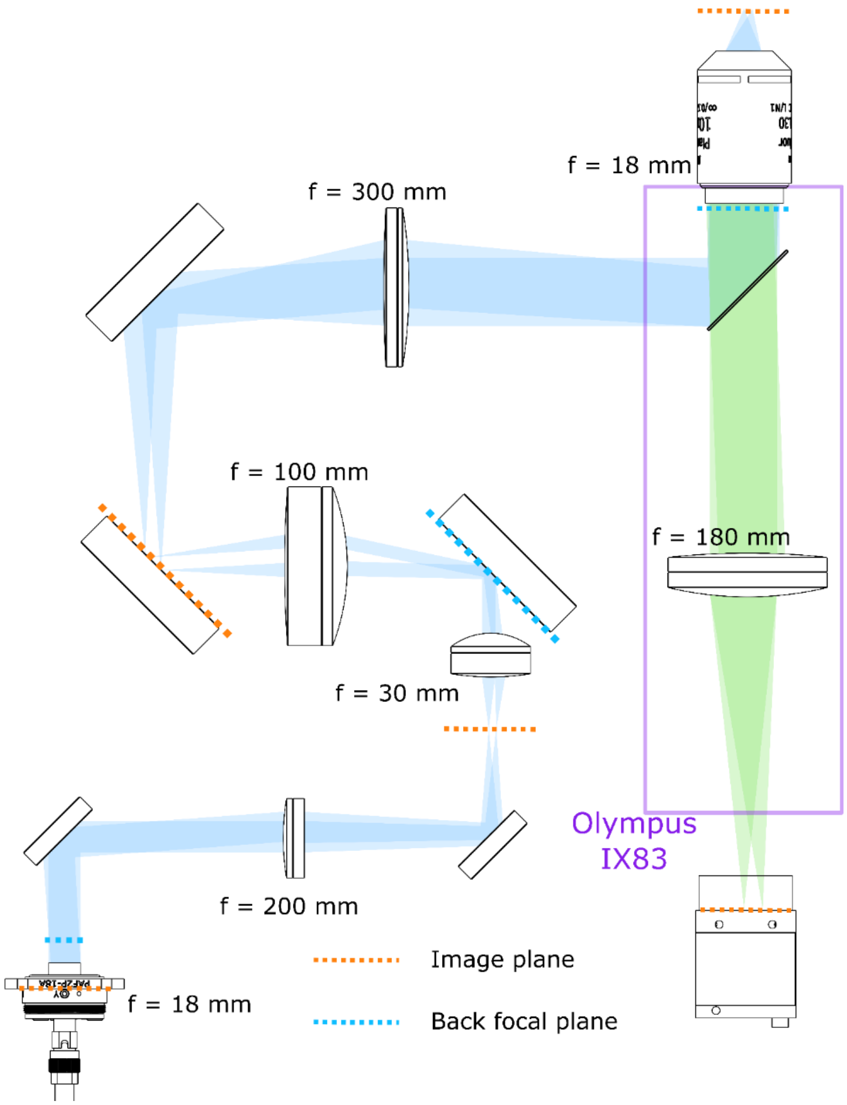


Figure S5 – Optical diagram of coupling of fiber output into Olympus IX83 microscope.

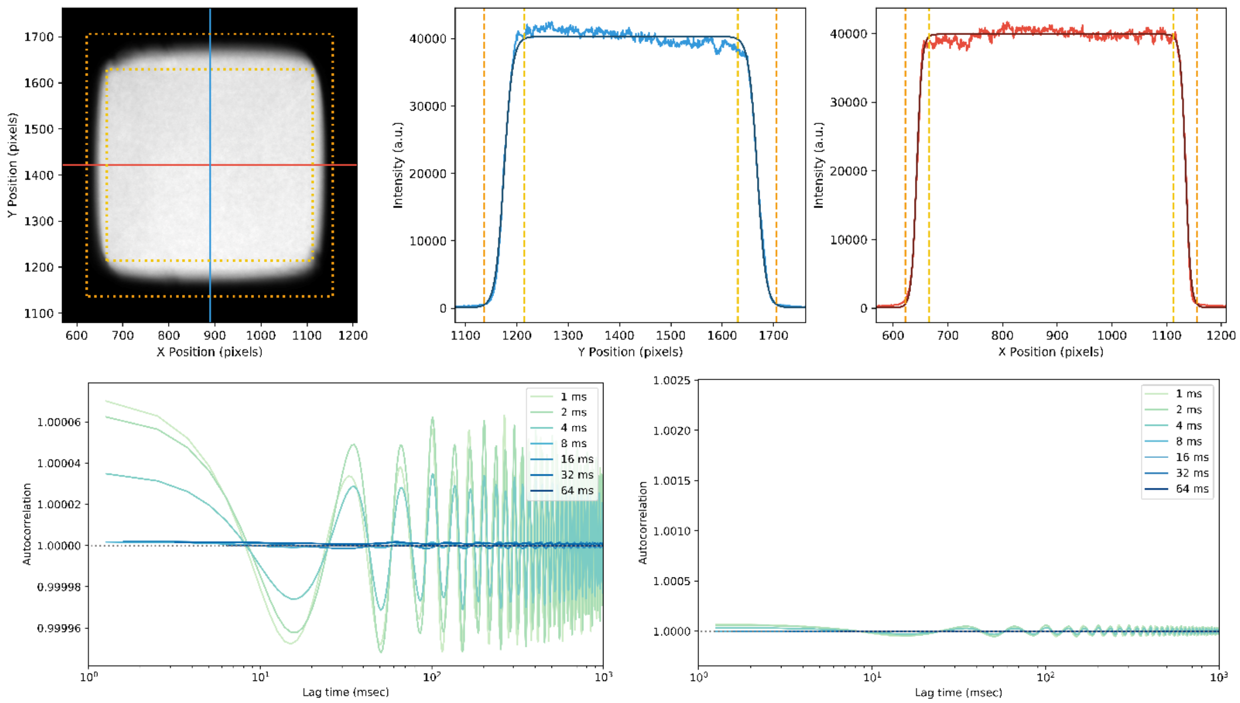


Figure S6 – Homogeneity images and autocorrelation traces from EvenField speckle reducer with Small Square fiber and FiberPort collimator at fiber exit. Top left panel shows fluorescence image of projected fiber face on test specimen. Top row, center and right - Blue and red lines indicate traces along X (red) and Y (blue) axes, with corresponding intensity profiles and fits in center and right columns. Dashed lines are 99% and 1% levels, defining the inner (yellow) and outer (orange) bounding boxes in intensity images. Bottom row shows autocorrelation trace at exposure times tested. Axes scaled to emphasize these data (left) and scaled to match that of circular fiber with Optotune speckle reducer (Figure 3 and Supplemental Figure 2).
